# Efficient Learning of Predictive Maps for Flexible Planning

**DOI:** 10.64898/2026.02.11.705395

**Authors:** Armin Bazarjani, Payam Piray

**Author notes:** Corresponding author: Armin Bazarjani, Payam Piray.

## Abstract

Cognitive maps enable flexible behavior by providing reusable internal representations of task structure. The successor representation, a predictive map that encodes expected future state occupancy, has been proposed as one way such maps might be computed in the brain, but its policy dependence severely limits flexible planning. We introduce a new model, the successor representation with importance sampling (SR-IS), which combines temporal-difference learning with importance sampling to construct policy-independent predictive maps. SR-IS learns the structure of the environment without being constrained by the agent’s current decision policy. These representations can be efficiently updated when the environment changes, enabling rapid behavioral adaptation. We show that SR-IS outperforms existing models in planning tasks and provides a better account of the graded biases in human replanning that previous models could not explain. This work bridges theories of predictive maps with observed planning behavior and offers new insights into flexible decision-making in the brain.

## Introduction

The ability to flexibly plan and adapt to changing environments is a fundamental aspect of intelligent behavior, and understanding its underlying mechanisms remains a central goal in neuroscience and psychology ^1–5^. It has been proposed that this ability relies on the brain’s computational representations, known as cognitive maps, which organize task-related information in a flexible and efficient manner ^3;6–9^. However, it is quite challenging to efficiently create representations that are shaped by our current goals and environment, yet also remain useful in the future. Despite numerous attempts, particularly those built on reinforcement learning (RL), existing computational models of planning face significant challenges. These models often struggle to achieve a balance between flexibility, efficiency in decision-making, and efficiency in updating their representations. Here, we introduce a new model to address these issues.

RL offers crucial insight into the brain’s capacity for efficient and flexible behavior, stemming from its ability to reuse previous computations, a concept that aligns with cognitive maps ^10;11^. However, a significant challenge lies in organizing these computations to adapt to new tasks, goals, or environmental changes, as the efficiency of reuse often conflicts with the need for flexibility ^12–14^. Consider this scenario: You leave your apartment for a morning coffee, navigating the city while simultaneously learning and updating your mental map of it. The next day, your destination changes to the library. Ideally, your cognitive map should support this new goal without being constrained by yesterday’s coffee shop route. While humans and animals may sometimes exhibit habitual behavior (like reflexively heading towards the coffee shop when stepping out of your house), they often demonstrate goal-directed or “model-based” behaviors ^15–20^. These behaviors efficiently utilize mental representations of tasks and environments to achieve current objectives. The key challenge lies in understanding how these flexible yet efficient representations are constructed and updated.

The successor representation (SR) ^21^, a prominent proposal from RL, attempts to address this challenge by caching expectations about future state visits ^22–25^. Importantly, the SR can be efficiently learned using the same “temporal-difference” (TD) ^26^ algorithms that are popular in RL and have been influential in understanding the role of phasic dopamine responses as prediction error signals representing the difference between observations and expectations ^27–34^. In the case of SR, the TD prediction errors represent the difference between the observed and expected future state visits. However, while the SR can be efficiently learned and updated, it struggles with flexibility. This inflexibility stems from the SR’s dependence on the specific policy used during learning—as the stored predictions reflect the agent’s past behavior rather than the underlying structure of the environment. Its reliance on old decision policies limits its usefulness when goals or environmental transitions change, resulting in behavior that is far more rigid than that observed in humans and animals ^24^.

Building on theoretical advances from control theory ^35;36^, a new RL model, linear RL constructs a computational map similar to the SR, but under a default policy ^37^. This policy is independent of previous or current goals, and the resulting map, called the default representation (DR), depicts the probability of visiting each future state when following this default policy. When the default policy is unbiased (e.g., uniform), the linear RL approach provides a highly accurate approximation of the optimal solution to RL problems, particularly those with deterministic transitions and specific goals ^37^. The solution offers two key advantages: First, it is a closed-form, linear, solution that does not depend on future values or actions, allowing for efficient implementation with a single layer of a neural network. Second, it enables efficient computation of map updates when transitions change or new goals are introduced. In principle, these updates affect the entire map, which can be as large as a matrix with dimensions equal to the number of states in the environment. However, they can be computed efficiently by leveraging a much smaller, low-dimensional representation whose size depends only on the scope of the changes in the environment. Yet, unlike the SR, there is no efficient TD algorithm for learning the DR. In this work, we address this issue.

Here, we introduce SR-IS (Successor Representation with Importance Sampling), a new model that leverages the principles of importance sampling from probability theory. In our context, this technique allows us to compute an unbiased, general successor map similar to the representation in linear RL, while the world is experienced under a specific goal-directed decision policy. Consider the coffee shop example again. As you leave your house to get coffee, you experience the successor states based on your current decision policy, which is shaped by your goal to get to the coffee shop. However, the computations that you ideally want to cache are the expected occupancies under a default policy (e.g., an unbiased uniform policy) that is independent of your current goal and reusable across future situations. Importance sampling enables us to do that.

We demonstrate that importance sampling can be effectively combined with the TD method to produce unbiased successor maps that are useful beyond the current goal and task. These maps represent likely future states under a default policy, despite being constructed from experiences derived based on a goal-directed decision policy. The solution is straightforward and closely resembles the classical TD algorithm for learning the SR. When the default and decision policies are matched, it reduces to the standard SR; otherwise, it smoothly debiases the SR. We show that such maps can be efficiently used within the linear RL framework, maintaining its advantages while also addressing its shortcomings. Importantly, the model explains aspects of human replanning behavior that earlier models struggled to account for ^25^, providing a parsimonious and normative explanation for graded differences and biases in human decision making.

## Results

### Model

#### Successor representation

In Markov decision tasks (such as navigating mazes or playing video games), an agent moves through a sequence of states. At each state, the agent receives a reward or punishment and must choose from available actions which determine the next state. RL captures this process by maximizing the expected sum of future rewards, called the “value” function.

Finding the optimal value function requires solving a series of interdependent optimizations, as the standard RL objective has no closed-form solution. Since this is often intractable, one simplified approach instead assumes a fixed decision policy *π* and calculate the value function under that policy, *V* ^*π*^(*s*). This represents the expected sum of (temporally discounted) future rewards from state *s* when following policy *π*. The SR emerges naturally from this framework, as *V* ^*π*^(*s*) can be calculated using the expected future occupancy of each state *s*′ along trajectories that start in state *s* and follow policy *π* ^21;23^.

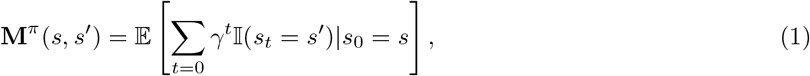

where *s*_*t*_ is the state visited at time *t, γ* ∈ [0, 1] is the discount factor parameter that downweights distal rewards, and I (*s*_*t*_ = *s*′) is an indicator function that is 1 if *s*_*t*_ = *s*′ and 0 otherwise. The superscript (*π*) in **M**^*π*^ indicates that the expectation is taken with respect to the decision policy distribution, *π*. Thus, two physically adjacent states that predict divergent future states under the decision policy will have dissimilar representations, and two states that predict similar future states will have similar representations. The SR is typically represented as a matrix **M**^*π*^, where **M**^*π*^(*s, s*′) is the element at position (*s, s*′).

An online TD learning approach has long been used to learn the SR, **M**^*π*^, from experiences. This approach caches rows of **M**^*π*^ and incrementally updates them after transitioning to a new state ^21^. Specifically, following a transition, the corresponding row of the state in which the transition was made from is updated as follows,

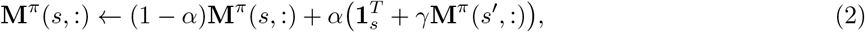

where **M**^*π*^(*s*, :) is the *s*th row of the SR matrix, 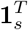 is a binary row vector that is all zeros except for a 1 in the *s*th position, and *α* is the step size parameter that controls the rate of learning. Although this may not be immediately obvious from Equation 2, one can see that the TD update for the SR puts more weight on the neighboring states (*s*′) that are visited more frequently from the current state (*s*). This is specifically where the policy-dependence of the SR comes from as it is a representation for the expected occupancy of states, thus if a state is visited more frequently under policy *π* the expected occupancy of that state should be higher.

#### Linear RL

A new model “linear RL,” dramatically simplifies RL problems by providing a closed-form solution to the value function ^37^, building on theoretical work from control theory ^35;36^. This approach approximates the original problem by introducing a new gain function for each state, defined as that state’s reward minus a control cost term. This control cost reflects the dissimilarity between the decision policy and a default policy, *π*^*d*^. Importantly, when the default policy is unbiased (e.g., uniform), the control cost term promotes stochastic behavior (traditionally useful for solving the explore-exploit dilemma) without biasing the resulting behavioral policy. As shown by Piray and Daw ^37^. this solution depends on a predictive map called the DR that is similar to the SR, but crucially it is constructed with respect to the default policy and not the decision policy. This addresses a key deficiency of the SR, which depends on a fixed (and typically outdated) policy.

However, learning the DR online presents significant challenges. In any online RL problem, the agent must act according to the current decision policy, not a uniform (i.e., random) default policy. Yet, using these actions directly with the TD rule would result in learning the map under the decision policy rather than the desired default policy. The key question becomes: how can we act according to the current decision policy while learning representations under the default policy that will remain useful even after the current policy becomes irrelevant?

#### Importance sampling

Importance sampling is a statistical method that enables efficient estimation of properties from a target probability distribution while sampling from an alternative distribution ^38;39^. It addresses a common challenge in statistics—calculating the expected value of a function under probability distribution *p* when we cannot solve the expectation analytically or sample directly from *p*. Simply drawing samples from a different distribution *q* would create a biased estimator, with the bias increasing the more *q* diverges from *p*. Importance sampling offers an elegant solution by drawing samples from the alternative distribution *q* (typically one that is easy to sample from) and correcting for the bias through “importance weights,” which are calculated as the ratio *p/q*. These weights adjust for sampling discrepancy by increasing the influence of samples that are likely under *p* but unlikely under *q*, while decreasing the influence of samples that are unlikely under *p* but likely under *q*. This approach constructs an unbiased estimator of the expectation.

#### New model: SR with importance sampling (SR-IS)

Let’s now return to the fundamental challenge in representation learning: how can we act according to our current policy while building representations that will remain useful when that policy becomes irrelevant? Importance sampling offers a solution. While we act according to the decision policy, we can learn the predictive representation for the default policy by applying importance weights, yielding an unbiased estimator of the DR (i.e., the SR under the default policy). We refer to this approach as SR-IS, which modifies the traditional TD learning algorithm:

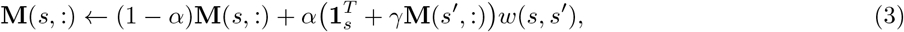

where 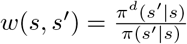 represents the importance weight, with *π*(*s*′|*s*) denoting the probability of transitioning from state *s* to state *s*′ under the decision policy *π*, and *π*^*d*^(*s*′ *s*), the corresponding probability under the default policy. This assumes a deterministic environment, where the next state is fully determined by the current state and action. The superscript *π* is omitted because matrix **M** is effectively learned under the default policy, which we assume to be uniform throughout this work (though the framework generalizes to any default policy). In this setup, the importance weights *w*(*s, s*′) correct for the discrepancy between the sampling distribution (decision policy *π*) and the target distribution (default policy *π*^*d*^). Specifically, transitions that occur more frequently under *π*^*d*^ than *π* receive higher weights, while transitions that are more probable under *π* than *π*^*d*^ receive lower weights. This reweighting mechanism ensures unbiased estimation of the default policy’s map, despite experiencing state transitions generated by a potentially very different decision policy.

To illustrate the key mechanisms of the model, consider a simple environment where an agent faces a choice between two reward states equidistant from its current location (Figure 1**a**). Although both states are equally accessible under a uniform default policy, they offer different rewards, with the right state providing a larger reward. The agent’s decision policy therefore favors movement toward the higher reward whenever it reaches this junction. Because the SR is policy-dependent, this behavioral preference alters its representation: the connection between the agent’s current state and the higher-reward state to the right is updated more frequently than the connection to the left state. Over time, the classical SR comes to overrepresent the more frequently visited state, even though both are equally reachable in spatial terms.

**Figure 1:**
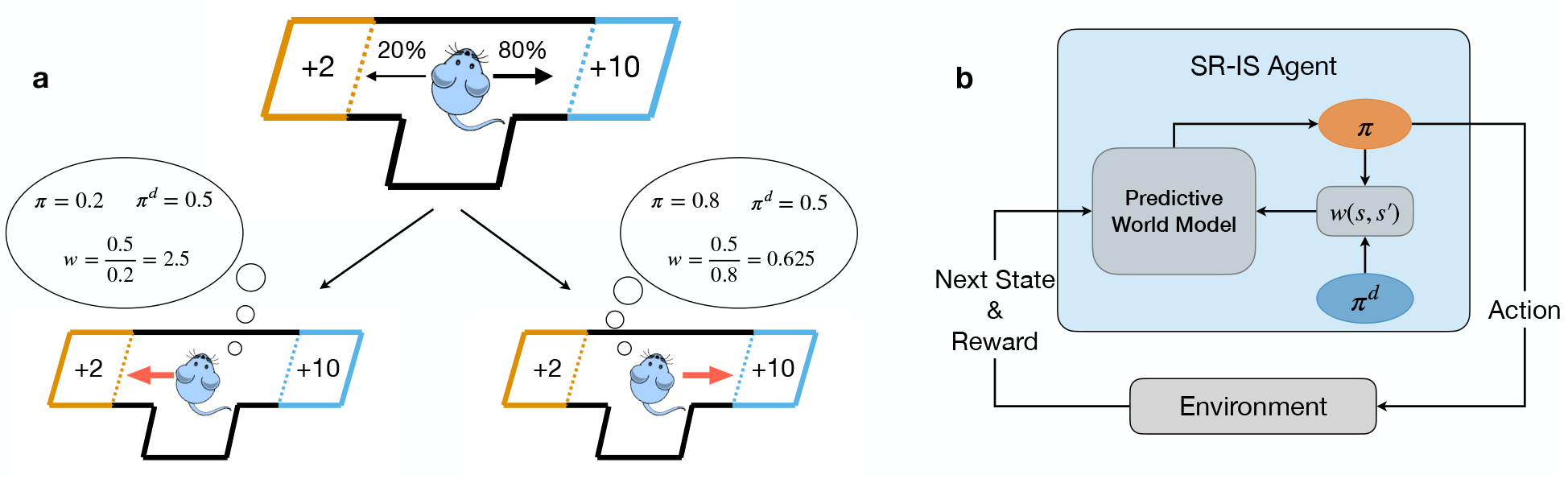
The SR-IS model. (**a**) Mechanism by which importance sampling debiases learned representations. After training, the agent preferentially moves right toward the higher reward, whereas the default policy remains constant and uniform. Because rightward movement is more probable under the decision policy, the resulting importance sampling term, *w*(*s, s*′), is smaller than if the agent chose the less probable leftward action. (**b**) High-level overview of the framework. We assume a classic RL setting where the SR-IS agent interacts with an environment. The interaction can be formalized with the agent acting on the environment with each action resulting in a new state and reward. The agent’s action is used to inform the importance sampling term which is then used, in conjunction with the classic TD update, to update the agent’s internal state-state representation.

This is precisely the kind of bias that the importance sampling term *w*(*s, s*′) is designed to correct. Intuitively, the importance sampling term *w*(*s, s*′) up-weights transitions that are more likely under the default policy *π*^*d*^ and down-weights those that are more likely under the decision policy *π* (Figure 1**a**). By incorporating *w*(*s, s*′) into the TD update rule (Equation 3), the model is able to de-bias its learned policy from the decision policy (Figure 1**b**). This allows the model to learn a representation that is less biased towards the specific actions taken by the agent during training and instead captures the underlying dynamics of the environment. As a result, the model learns a more robust and adaptable cognitive map that can support flexible planning and generalization to new tasks.

## Model Performance

### SR-IS converges to the DR

We first validate our approach by showing that the representation learned by SR-IS converges to the unrealistic DR matrix obtained via matrix inversion. This is theoretically important because SR-IS is designed to learn the DR map, a representation that enables linear RL methods, without requiring complete environment knowledge. We refer to the DR obtained through matrix inversion as the “Complete” model, because it uses perfect, complete, knowledge of the environment’s transition structure to compute the DR. In the limit, the representation learned by SR-IS should converge to this map.

To test convergence, we used a classic RL benchmark ^40^, the four-room environment (Figure 2**a**,**b**). In this environment, we compared three key quantities: the representation learned by SR-IS through online temporal difference learning with importance sampling, the representation learned by the standard SR without importance sampling, and the Complete DR computed directly through matrix inversion. We measured the mean absolute difference between these representations as the two agents (SR, SR-IS) navigated the environment and updated their respective representations.

**Figure 2:**
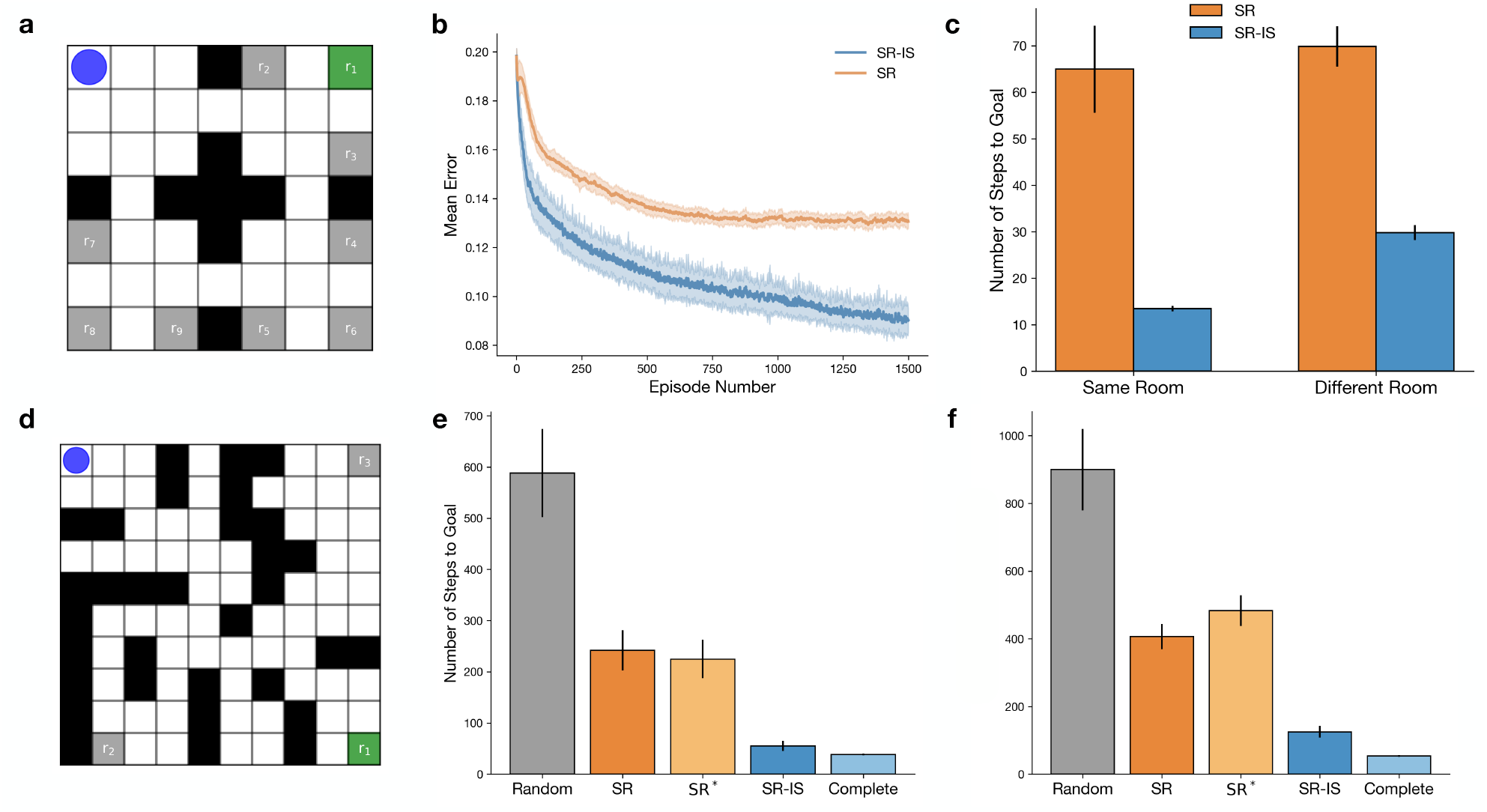
SR-IS enables flexible and efficient replanning. (**a**) Four-room environment showing the agent’s starting location (blue circle) and the initial reward state r_1_ (green square). (**b**) The representation learned by the SR-IS converges to that of the Complete model (which uses unrealistic matrix inversion to compute the DR map), in contrast to that of the SR model (the same model without importance sampling). The plot shows the mean absolute difference between each model and the Complete model during navigation to r_1_. (**c**) Replanning comparison between SR and SR-IS models: after learning representation with respect to r_1_, agents replan to 8 new terminal states, r_2*−*9_, with 2 in the same room as r_1_ and the other 6 evenly distributed in other rooms. SR-IS maintains efficiency across rooms while SR only replans effectively within the same room. (**d**) A relatively complex maze showing the agent’s initial position (blue circle) and the initial goal state r_1_ (green square). Unlike the previous simulations, r_2_ and r_3_ were not initially designated as terminal states, requiring the use of Equation 4 when replanning. (**e**) Performance when replanning to r_2_ after initial training to r_1_: SR performs near the random-walk level, SR* (SR combined with Equation 4) remains biased toward the previous reward, and SR-IS nearly matches the Complete model’s performance. (**f**) Same as (**e**), but for replanning to r_3_. Error bars represent the standard error of the mean. See Supplementary Fig. 1 to see validation of SR-IS on two classical replanning paradigms introduced by Tolman.

The results revealed that SR-IS converges to the Complete DR as the number of learning episodes increases (Figure 2**b**), with the difference between the two representations approaching zero. This confirms that importance sampling enables the model to learn the correct underlying structure through experience alone, without requiring expensive matrix operations. In contrast, the standard SR without importance sampling converges to a distinctly different representation that reflects the specific policy used during training.

### SR-IS demonstrates cross-room planning efficiency

We next evaluated how the unbiased nature of the map learned by SR-IS affects planning flexibility across different spatial domains (Figure 2**c**). After training both SR-IS and SR models to navigate to an initial goal state (r_1_) in the four-room environment, we tested their ability to replan routes to eight new goal states (r_2_, …, r_9_). In these experiments, each goal state was treated as a terminal state, with the only manipulation being the reward value at each terminal location. Both models were initially trained with r_1_ as the sole rewarding terminal state, producing a representation of the environment that reflected that configuration. We then used these learned representations to assess each model’s capacity for replanning to the remaining eight goal states. Critically, no retraining was performed after the reward structure was changed; Instead, we directly used the representation learned during the training phase to derive new policies for the alternative goal locations.

SR-IS demonstrated consistent performance across all goals, efficiently computing paths regardless of the goal’s location relative to r_1_. In contrast, the SR model showed a strong spatial bias. While it maintained effectiveness for goals within the same room as r_1_, its performance degraded substantially for goals in other rooms. This disparity reflects the SR’s policy dependence, which arises because it builds a representation biased toward frequently visited states under the r_1_-directed policy, often first going to the room in which r_1_ is located, resulting in suboptimal routes to new terminal locations. SR-IS overcomes this limitation through importance sampling, learning representations under the default policy that remain useful when the goal changes.

These results complement our convergence findings by demonstrating that SR-IS’s theoretical advantages translate into practical benefits for spatial navigation and planning. The model’s ability to maintain performance across spatially distinct regions highlights the importance of learning unbiased representations for flexible behavior.

### The representation learned by SR-IS is able to replan with a single update

As shown by Piray and Daw ^37^, one of the most important features of the DR map used by the linear RL framework is that it can be efficiently and exactly updated when state transitions change. This capability builds on matrix algebra (specifically the Woodbury matrix inversion lemma ^41;42^), which allows us to update the representation of **M** in place by accounting for local changes in the transition graph:

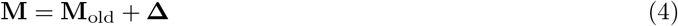

where **Δ** is a low-rank, easy-to-compute change matrix whose rank equals the number of states with modified transitions (see Methods for the exact definition). This is important for adapting both to structural changes in the environment, such as new barriers or cases when previous non-goal states become goals, or vice versa. Since SR-IS converges to the same DR Complete map, it can leverage the same efficient update method, inheriting all the computational advantages of the linear RL framework.

We conducted simulations in a more complex maze environment to demonstrate this point. Similar to the previous simulation, we tasked the model with replanning to new terminal states. The crucial difference here is that the model lacked prior knowledge of which states were terminal. Thus, each time a new state became terminal, the transition structure of the environment changed, requiring an update to the learned map. We first used the SR-IS model to learn the map while the agent planned a route from a starting state in one corner to a goal state in the opposite corner of the maze, (r_1_). We then tested whether this learned representation, combined with Equation 4, could find the shortest path to two non-goal states, r_2_ and r_3_ (Figure 2**d**).

Simultaneously, we compared SR-IS against a number of alternative models in the same maze, including the Complete model (i.e., using matrix inversion to calculate the DR map) and the standard SR model. Each model first constructed a predictive map with an initial goal state r_1_ (Figure 2**d**). We then changed the goal to r_2_ or r_3_ and evaluated how well each model could reuse its previous map to plan a new path. SR-IS, on average, was the closest match to the Complete model (Figure 2**e**,**f**), a notable achievement given that the Complete model relies on a computationally intensive matrix inversion that is infeasible for biological systems.

In contrast, the SR model’s behavior was closer to a random walk for both new goal locations. This is expected, as SR’s policy dependence causes its representation to change substantially with the introduction of new goals. For comparison, we also tested a second version of SR, labeled SR*, which used the same update method as SR-IS and the Complete models (Equation 4). While one might expect this approach to help, it was mathematically clear from the outset that it would not benefit the SR. Equation 4 is only effective when the change produces a local alteration in the transition structure of the learned map. For SR-IS and the Complete model, this locality holds because a new goal only changes the default policy for successor states of the new and old terminal states. In contrast, for the SR, the entire transition structure must change, since its map is tied to the decision policy, which itself changes completely with the introduction of new goals. Our analysis revealed that SR* was only marginally better than the SR.

### SR-IS replans similarly to humans

Next, we tested the model on a task introduced by Russek et al. ^24^, known as policy revaluation, to highlight the SR’s key limitation (Figure 3). They showed that multiple implementations of SR, including the unrealistic but most powerful version that computes the SR matrix via direct inversion, fail to solve this task. This failure stems from SR’s inherent policy dependence, which prevents it from adapting when a complete policy revaluation is required. In contrast, the Complete linear RL model solves the task almost perfectly, consistent with its close correspondence to model-based behavior. SR-IS also succeeds, though with a slight SR-like bias, a characteristic signature of the model. This reduction in performance arises from high sampling variance when states common under the default policy are rarely visited by the decision policy. The effect is most evident when the environment is undersampled, presumably because the matrix learned by SR-IS has not yet converged to the complete DR (Supplementary Fig 2). This reflects a fundamental limitation of importance sampling that, while unbiased in the limit, exhibits high variance when the sampling distribution poorly covers regions important to the target distribution.

**Figure 3:**
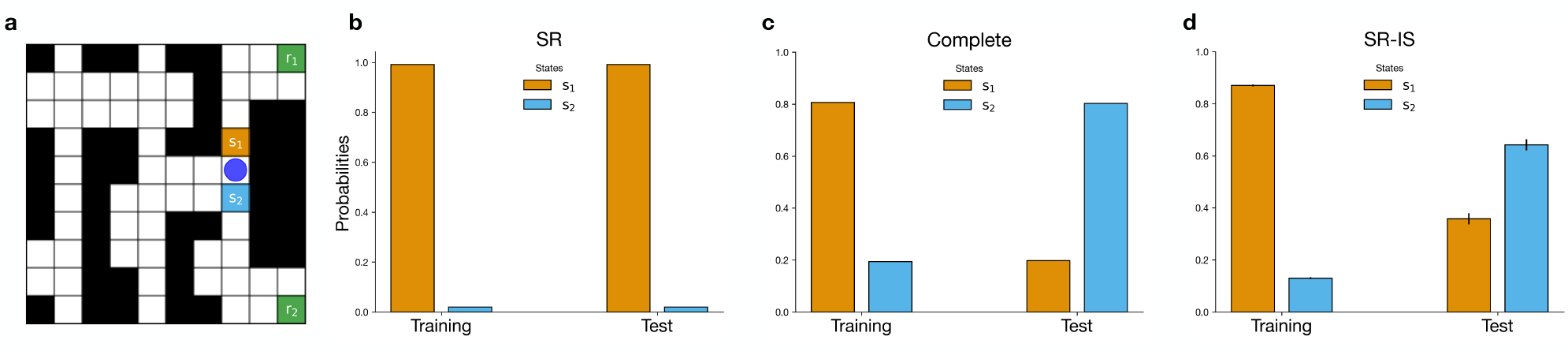
SR-IS solves revaluation tasks that defeat SR. (**a**) Policy revaluation task from Russek et al. ^24^. The maze shows the agent’s location with a blue circle and two neighboring states (s_1_ & s_2_) showing different paths to the two opposing terminal states (r_1_ & r_2_). (**b**-**d**) The probability of the SR (simulated here with direct matrix inversion, which is the most powerful implementation of the SR), Complete, and SR-IS agents to select either state s_1_ or s_2_, effectively showing the agent’s preferences over both terminal reward states r_1_ and r_2_ during training and testing. (**b**) The SR agent struggles from policy-dependence and is therefore unable to effectively navigate to the higher reward (r_2_) during test. (**c**) The Complete model (linear RL) is able to perfectly update its representation in correspondence to the higher reward. (**d**) The SR-IS agent constructs a policy-independent representation and is therefore able to change preference upon learning of the higher reward albeit with a slight reduction in performance due to sampling variance. Error bars represent the standard error of the mean. See Supplementary Fig. 2 to see how SR-IS’s bias becomes less salient as the number of training steps increases.

This limitation, rooted in SR’s policy dependence, raises a broader question: how do humans handle situations that require substantial changes to an existing policy? To address this, we next examine a set of replanning tasks that reveal intriguing but puzzling patterns of both flexibility and inflexibility in human behavior. Built on the insights from Russek et al. ^24^ (Figure 3) and tested experimentally by Momennejad et al. ^25^, these tasks provide an ideal testbed for evaluating our model against existing alternatives. The experimental paradigm consists of three cleverly designed tasks that probe specific aspects of revaluation behavior. A key finding was that humans display an asymmetric pattern of flexibility across these replanning tasks, a pattern that could not be explained by either the SR model or model-based accounts (such as the Complete linear RL model shown in Figure 3).

The three tasks, termed reward, policy, and transition revaluation, are illustrated in Figure 4**a**. In these experiments, human subjects were initially trained to navigate a three-stage sequential task leading to one of three terminal states. The training phase was followed by a revaluation phase, during which participants either experienced a significant reward change in a previously disfavored terminal state or learned about a change in the transition structure. Importantly, they did not have to go through the entire task again to experience this change. In the final testing phase, the participants were placed back at the beginning of the maze.

**Figure 4:**
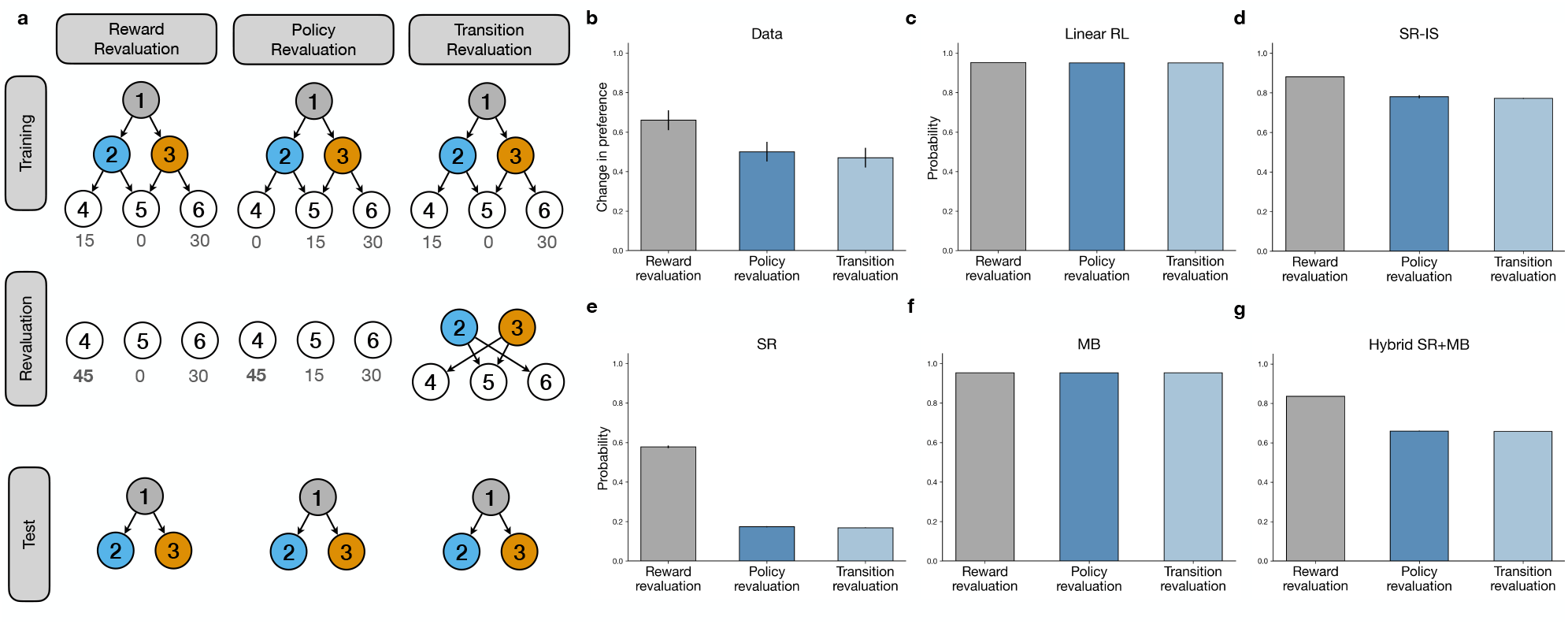
SR-IS shows human-like planning behavior. (**a**) The underlying structure of the reward, policy, and transition revaluation experiments in Momennejad et al. ^25^ (**b**) Reconstruction of human data from the task ^25^, more participants changed their preference (policy) during reward revaluation than policy revaluation. (**c**-**g**) We compare the human data with model performance showing the model’s probability of switching states for the linear RL, SR-IS, SR, model-based (MB), and hybrid SR+MB models. The SR-IS model accurately recapitulated the asymmetry in preference switching in the reward and policy revaluation conditions, producing results closely matching those of the descriptive hybrid model. Error bars represent the standard error of the mean. See Supplementary Fig. 3 for full performance of SR-IS model during training and testing.

For the policy and transition revaluation tasks, the new optimal action requires a change in preference that is directly antithetical to the current policy. Even though the reward revaluation setting also requires humans to make a change in preference, the path from state 1 to state 2 was still rewarding and allowed the participant to still achieve the maximal reward, whereas in the policy and transition revaluation setting going to state 2 was never as rewarding as state 3. The SR model, which is learned with respect to the decision policy used during the training phase, cannot adapt in the test phase for these settings.

The empirical results ^25^ showed that participants were generally able to modify their behavioral policy despite never having directly reached the new terminal state through actions in the maze. Sensitivity, measured as the change in preference for choosing state 2 from state 1 after revaluation, revealed a key asymmetry: participants showed significantly higher sensitivity in the reward revaluation condition compared to both the policy and transition revaluation conditions (Figure 4**b**).

Neither the standard SR model, nor the Complete linear RL model and model-based models can adequately explain this pattern of behavior (Figure 4**c**,**e**,**f**). The standard SR model, being inherently policy-dependent, is expected to entirely fail at policy revaluation as it cannot update its cached predictions without direct experience under the new policy ^24^. Conversely, linear RL and model-based models, with their perfect knowledge, predict equally good performance across all three conditions, failing to capture the graded sensitivity differences observed in human behavior.

The original study interpreted this behavioral asymmetry as evidence for the SR, noting that a hybrid model combining the SR with model-based RL could reproduce the observed pattern (Figure 4**g**). While such a hybrid model provides a good fit to the data, it offers no clear computational rationale for why the brain would maintain two redundant planning systems—a point that is particularly problematic given that the SR was originally proposed as a computationally efficient alternative to model-based RL. The hybrid approach paradoxically reintroduces model-based costs at decision time while retaining the SR’s inherent inflexibility, all without specifying a formal mechanism for system arbitration. While the brain may indeed arbitrate between distinct systems ^17^, such mechanisms typically invoke domain-general principles ^43^ like uncertainty-based switching ^5^ or speed-accuracy tradeoffs ^16^. In contrast, the existing hybrid model fails to explain why this behavior emerges. In effect, the hybrid model stitches together components with incompatible assumptions; it takes a descriptive approach that fits behavioral asymmetries through parameter mixing rather than a normative one that explains them through fundamental computational principles.

Simulating SR-IS on this task yields a normative account of the observed “hybrid pattern” (Figure 4**d**). Its success arises from a key insight: the same importance sampling mechanism that enables flexible replanning also naturally produces human-like limitations, especially in policy and transition revaluation tasks. As a result, some simulated agents, like their human counterparts, fail to make optimal choices. This distinctive combination of successful replanning with graded performance differences emerges not from parameter tuning or mixing systems, but from a fundamental property of importance sampling whereby limited experience with rarely visited states leads to variability and occasional suboptimality. SR-IS therefore provides a normative explanation for both human competence and human limitations within a single computational framework.

To further substantiate the ability of the SR-IS model to capture seemingly dual-process behavior, we simulated our model on a task by Kahn and Daw ^44^. Participants performed a two-step decision task with alternating trial types. In choice trials (Figure 5**a**), they selected between two first-stage options, then between two second-stage options, receiving binary rewards. In observation trials (Figure 5**b**), they passively observed second-stage outcomes without making choices. This design tested how reward information gained through observation (without action) influenced subsequent choices.

**Figure 5:**
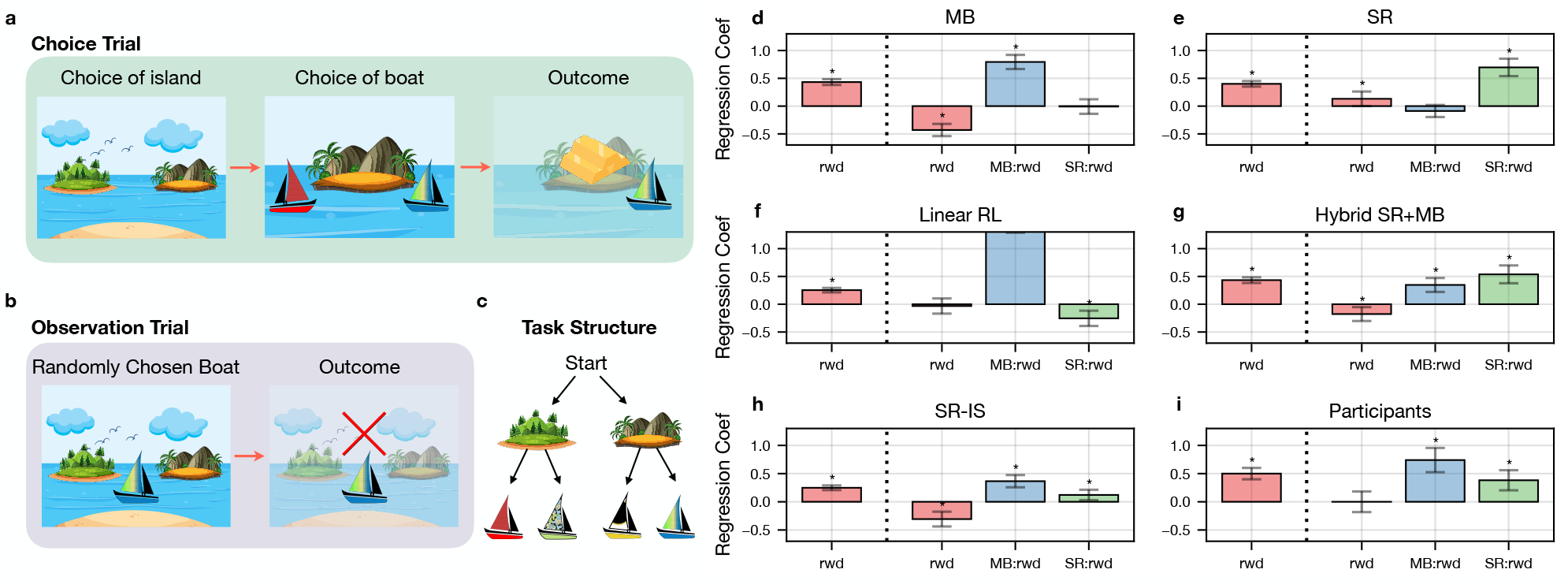
SR-IS shows human-like planning behavior. (**a, b**) The experimental procedure of Kahn and Daw ^44^. Trials alternate between “choice” trials where subjects choose both an island and then a boat, and “observation” trials where a randomly selected boat and its reward is delivered to the subject, importantly without any island choice or presentation. (**c**) Full schematic of trial structure showing the start state, choice of islands, and choice of boats at each island. (**d**-**h**) Analyses by the authors highlight key factors influencing how observation trials affect subsequent choices. The leftmost bar shows results from the reward-only model, where all four agent types demonstrate significant modulation by reward from the previous trial. The remaining three bars show results when model-based (MB) and successor representation (SR) interaction terms are included. MB signature: Because the MB agent maximizes over subsequent boat rewards (*r*_1_ and *r*_2_) at each island, 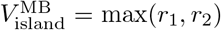, it is sensitive to whether either boat is rewarded. SR signature: Because the SR agent predicts values in the form of an expected occupancy, 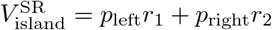, it is sensitive to its previous choices at that island (due to policy dependence). Both MB and Linear RL agents exhibit strong interactions with neighboring boat outcomes, while SR agents show strong interactions with prior choices at the sampled island. Both Hybrid SR+MB and SR-IS agents display both interaction effects with the Hybrid SR+MB agents emphasizing prior policy more heavily and SR-IS agents emphasizing the prior reward more heavily. Plots **d, e, g, i** are reproduced from the main paper. Error bars represent 95% CI. ^44^

The authors used a model-agnostic approach to analyze behavior by identifying unique trial-by-trial signatures that distinguish SR from model-based planning strategies. The model-based signature captures recursive maximization: a model-based agent compares boat values at each island, so observing a boat’s reward only influences subsequent choice if the neighboring boat has low value (i.e., when the comparison is decision-relevant). The SR signature captures on-policy state averaging: an SR agent predicts values from expected future occupancy, which depends on past choices, so observing a boat’s reward only influences subsequent choice if that boat was previously chosen (i.e., when it’s part of the learned trajectory). They quantified these signatures through interaction terms in a linear mixed model of choice behavior following observation trials. By leveraging these distinct behavioral markers, they could quantify each participant’s reliance on either strategy without assuming a specific computational model. Their analysis revealed that human participants employed both strategies, with the relative contribution of each varying across individuals and task conditions (Figure 5**i**). Importantly, neither pure model-based (Figure 5**d**) nor pure SR (Figure 5**e**) planning could fully account for the observed behavior. Instead, participants showed signatures of both recursive maximization (comparing current options) and on-policy averaging (learning from past choices), which was interpreted as requiring a hybrid account combining SR and model-based components (Figure 5**g**). However, while such a hybrid description can match behavior, it does not provide a normative explanation for why or when these two strategies should be combined. Simulations of pure model-based and pure SR accounts confirmed that each captured only its corresponding interaction term, whereas the Complete linear RL model (Figure 5**f**) behaved similarly to the model-based account as expected.

Simulating SR-IS on this task provides a normative account of the observed hybrid pattern. Like humans, it shows significant model-based and SR interaction terms (Figure 5**h**), arising from its unbiased approximation of the Complete linear RL model and residual SR-like effects from sampling variance, which are amplified when low-probability actions occur under the decision policy. Consistent with the replanning experiments from Momennejad et al. ^25^, SR-IS displays behavioral patterns characteristic of a mixture of SR and model-based accounts.

Beyond simulations, we fitted the SR-IS model to the data and performed a formal comparison against the SR, model-based, and hybrid accounts. The results indicate that SR-IS is the most likely model at the population level, with a protected exceedance probability ^45;46^ of 0.99 (Table 1).

**Table 1:** Model comparison results for Kahn and Daw ^44^.

| Model | Model Frequency | Protected Exceedance Probability |
| --- | --- | --- |
| SR | 0.06 | 0.00 |
| Model-based (MB) RL | 0.24 | 0.01 |
| Hybrid SR+MB | 0.25 | 0.01 |
| **SR-IS** | **0.45** | **0.99** |

### SR-IS results in a better match to rat and human navigation trajectories

The previous analyses demonstrated the advantages of the SR-IS model in experimental replanning paradigms. Here, we examine the model using the rich dataset from de Cothi et al., which stands out for both its comprehensive set of complex mazes and its more naturalistic experimental setup. de Cothi et al. ^47^ conducted a comprehensive cross-species comparison using their “Tartarus maze” - an innovative experimental paradigm requiring rapid adaptation to changing environmental obstacles while maintaining goal-directed navigation.

In their experiment, the goal state was fixed and then they had both humans and rats navigate through 25 different maze configurations, each having 10 different starting states. Rats were tested with a physical instantiation and humans were tested with immersive head-mounted virtual reality. Their findings revealed that both species showed remarkable similarity in their navigation patterns, with performance most closely matching SR agents compared to model-based or model-free accounts. However, they noted that the SR model struggled with certain maze configurations, particularly those requiring significant policy changes, a limitation we hypothesized could be addressed through SR-IS.

Our analyses confirmed this hypothesis. Following the original study ^47^, we fitted the SR-IS model and compared performance across trails as well as maze configurations. Across trials, we observed that the SR-IS model accounted better for 57% of the human trials and 52% of the rat trials (Figure 6**a**,**b**) Across maze configuration, we observed substantial variations with the SR-IS model showing an advantage across most maze configurations. Of course, these comparisons do not account for differences in number of parameters per model. Therefore, we also observed a formal model comparison we also conducted formal model comparison, which takes into account complexity (i.e., number of free parameters) of each model. SR-IS was the most frequently expressed model for both humans and rats (Table 2) with higher protected exceedance probability across all models.

**Table 2:** Model comparison results for de Cothi et al. ^47^.

| **Model comparison on humans** |  |  |
| --- | --- | --- |
| Model | Model Frequency | Protected Exceedance Probability |
| Model-free RL | 0.04 | 0.00 |
| Model-based (MB) RL | 0.04 | 0.00 |
| SR | 0.26 | 27.78 |
| Hybrid SR+MB | 0.04 | 0.00 |
| **SR-IS** | **0.61** | **72.22** |
| **Model comparison on rats** |  |  |
| Model | Model Frequency | Protected Exceedance Probability |
| Model-free RL | 0.07 | 0.00 |
| Model-based (MB) RL | 0.07 | 0.00 |
| SR | 0.14 | 11.11 |
| Hybrid SR+MB | 0.07 | 0.00 |
| **SR-IS** | **0.64** | **88.89** |

**Figure 6:**
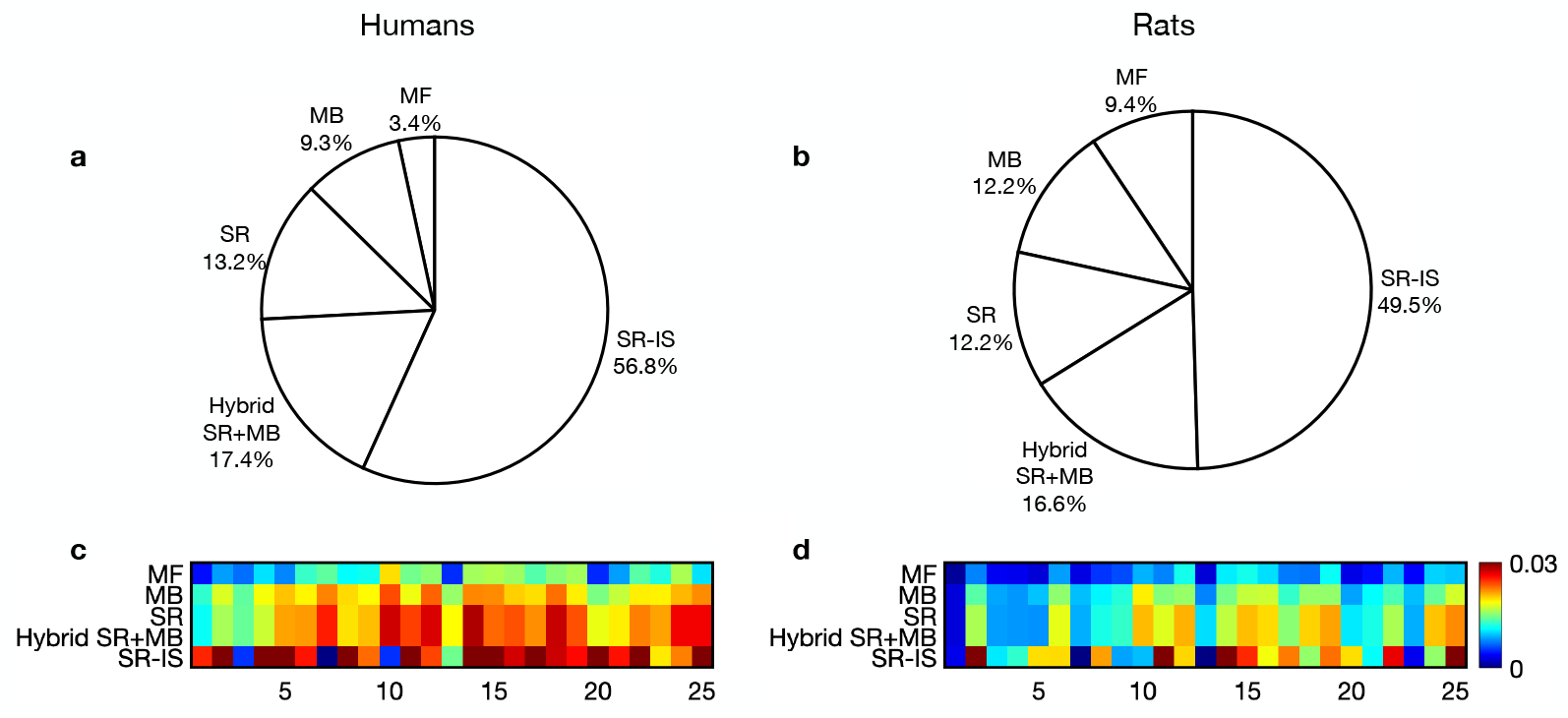
Maximum likelihood analyses across human and rat trajectories. (**a, b**) Proportion of trials best explained by the SR, SR-IS, and model-based (MB) models (maximum-likelihood assignment). The SR-IS agent was the maximum likelihood model to explain the behavior for the majority of trials. (**c, d**) Average likelihood per timestep compared to baseline across all maze configurations. We can see that the agent likelihoods vary across maze configurations for both humans and rats with the SR-IS agent generally having a stronger fit across all configurations. See also Supplementary Fig 4.

Further analysis also reveals a subtle but important aspect of SR’s policy-dependence that emerges in practice: non-uniform convergence due to experience-dependent learning. While SR theoretically converges to the same representation regardless of starting location, realistic learning scenarios with limited experience produce substantially different SR matrices depending on where learning began. This occurs because different regions of the environment receive unequal sampling during exploration under the behavioral policy. Consequently, SR’s performance becomes critically dependent on both the quantity and distribution of past experience, a direct consequence of its policy-dependent learning mechanism. We have highlighted two mazes that show this vulnerability more clearly because they contain critical decision points where optimal behavior requires well-learned representations of regions that may have been rarely visited under the original behavioral policy (Supplementary Fig. 4). These mazes are ideal for demonstrating this issue because the starting locations alternate between opposite sides—left/right in Maze 15 and top/bottom in Maze 22. While humans and rats readily adapt to these starting position changes once they understand the goal location and barrier structure, SR struggles because its policy-dependent learning prevents it from constructing a general representation of the environment layout.

SR-IS addresses this limitation, because it appropriately reweights experiences through importance sampling to ensure uniform learning across the state space. By correcting for the bias introduced by the behavioral policy during learning, SR-IS maintains a more consistent performance regardless of which regions were extensively explored during training. This advantage is particularly evident in the presented mazes, where SR-IS successfully navigates from novel starting positions that challenge standard SR.

### Distinctive predictions in stochastic environments

We next sought to identify situations where SR-IS makes distinctive predictions compared to alternative models. Following Piray and Daw ^37^, we consider a two-step decision task where one state leads to stochastic action outcomes (Figure 7**a**). We evaluate the models across two conditions: before and after reward revaluation. Before revaluation, SR-IS fails to adequately account for this stochasticity and incorrectly prefers state *S*_2_ when starting in *S*_1_, whereas SR, model-based, and hybrid all correctly prefer *S*_3_. This failure arises because SR-IS, like linear RL, assumes fully controllable transitions—an approximation that breaks down when stochasticity substantially affects the value of future states. Interestingly, after revaluation, the predictions of the models diverge further (Figure 7**b**–**e**). When terminal state rewards are changed to make the expected value of *S*_2_ higher than *S*_3_, SR-IS maintains its original (and now correct) preference. The SR, constrained by its policy dependence, continues to prefer *S*_3_ despite revaluation. Meanwhile, the model-based agent, with complete knowledge of the environment, fully reverses its preference. The hybrid model falls between the SR and model-based predictions, reflecting its mixture architecture. Note that these results also echo a previous computational account of model-based decision making that is a closely related formalism to linear RL ^48^. These contrasting predictions are empirically testable. SR-IS predicts that humans should either show greater errors in stochastic environments or avoid such errors by reverting to more costly iterative planning, measurable through longer response times. To our knowledge, these predictions remain untested and could distinguish between these competing computational accounts.

**Figure 7:**
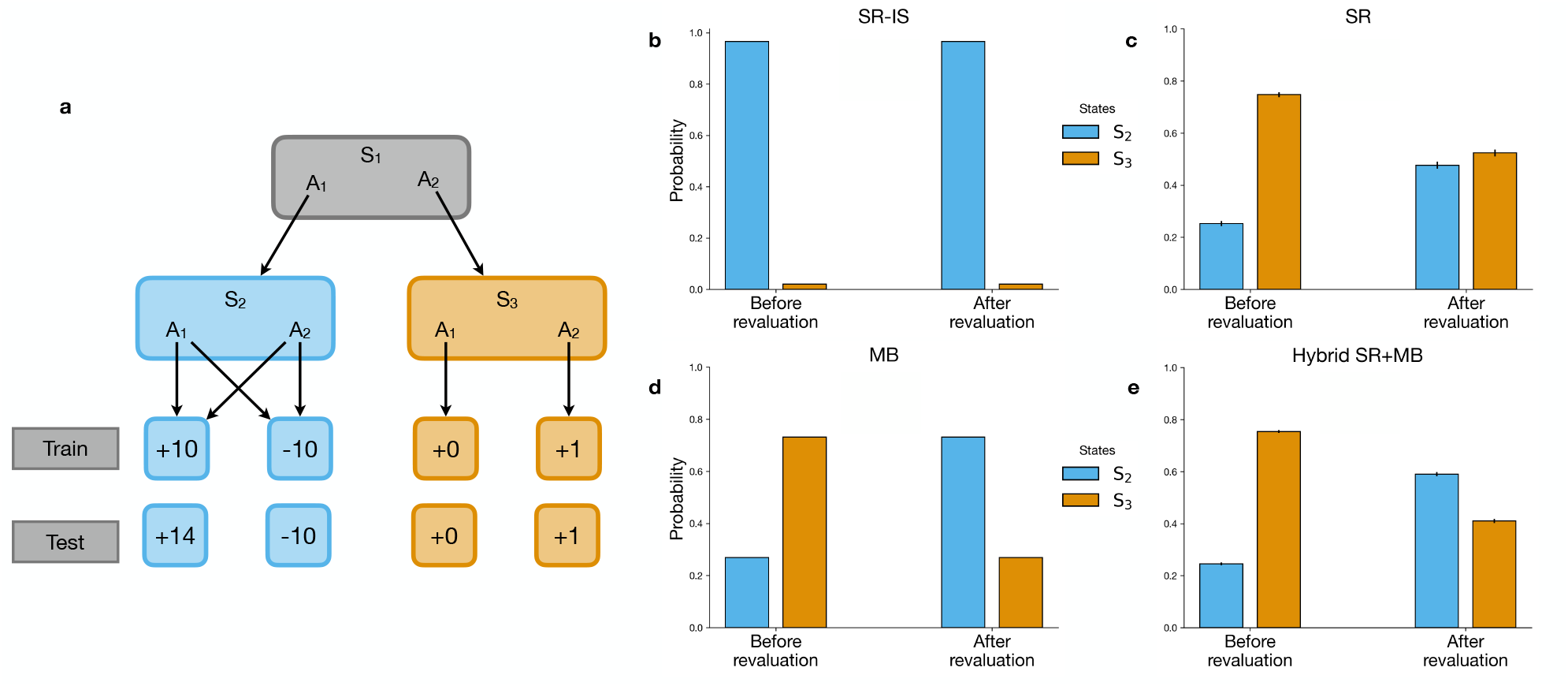
SR-IS in environments with stochastic transitions. (**a**) A task with stochastic transitions where SR-IS fails to exhibit optimal behavior. At state *S*_1_, actions *A*_1_ and *A*_2_ lead deterministically to *S*_2_ and *S*_3_, respectively. However, taking actions in state *S*_2_ leads stochastically to two different terminal states with +10 and −10 rewards (each with a 50% probability). Consequently, before revaluation, the expected value of state *S*_3_ is higher than that of *S*_2_, making *A*_2_ the optimal action in *S*_1_. After revaluation, the terminal state rewards change such that *A*_1_ becomes the optimal action. (**b–e**) Model preferences in state *S*_1_ (choosing between *S*_2_ and *S*_3_) both before and after the revaluation condition. SR-IS remains unchanged. The SR, due to its policy dependence, cannot adapt to the change. Model-based (MB) agent optimally switches preferences due to its complete knowledge of the transition structure. The Hybrid SR+MB model predicts a preference that falls intermediate to the SR and MB models.

## Discussion

The present work introduces SR-IS, a novel computational model that addresses a fundamental challenge in cognitive map construction: how to create task representations that are both flexibly reusable across contexts and computationally efficient through cached planning computations. This advances beyond previous RL approaches, which have been limited either by their reliance on specific policies (like SR) or by their dependence on computationally intensive processes (like model-based accounts ^5;17;24^ or matrix inversion ^37^).

At the core of SR-IS is a reweighting mechanism that systematically debiases learned representations. This mechanism effectively rebalances the learning process by increasing the influence of transitions that are more likely under the default policy while reducing the impact of those dictated by the current decision policy. By incorporating this weighting into the TD learning framework, the model systematically removes biases stemming from specific decision policies and instead constructs a representation that better reflects the environment’s inherent structure. This approach yields a more general and robust predictive map that maintains its utility across different tasks and goals, enabling flexible decision-making even in novel situations. Such an unbiased representation stands in contrast to traditional cached successor maps that are typically constrained by the specific experiences accumulated during training.

Our results demonstrate that SR-IS provides a unified computational account for behavioral patterns previously attributed to dual-system architectures. Across two distinct experimental paradigms, the revaluation tasks of Momennejad et al. ^25^ and the two-step decision task of Kahn and Daw ^44^, SR-IS reproduced the full spectrum of human behavior without invoking separate planning systems. In the revaluation experiments, SR-IS captured both the successful adaptation and the graded performance differences across task types, with policy and transition revaluation proving more challenging than reward revaluation. Similarly, in the two-step task of Kahn and Daw, SR-IS exhibited both model-based and SR signatures that characterize human choice patterns. Crucially, these diverse behavioral patterns emerge from a single computational principle: importance sampling enables flexible replanning while naturally introducing variance that manifests as human-like limitations. This variance is most pronounced when the decision policy rarely visits states that are common under the default policy, explaining why certain revaluation conditions prove more difficult. Rather than requiring ad hoc mixing of incompatible systems, SR-IS shows how apparent evidence for hybrid architectures may instead reflect the natural consequences of learning reusable representations through importance-weighted experience. This work thus advances our understanding of biological planning by demonstrating that behavioral flexibility and its limitations can arise from a single, normatively motivated computational mechanism.

The SR-IS model seamlessly integrates with the linear RL framework while addressing its primary limitation ^37^: the absence of an efficient learning algorithm. Through importance sampling, the model learns planning-ready representations even during goal-directed policy execution, similar to how individuals construct a general cognitive map of their city while traveling specific routes. When paired with an unbiased default policy, this approach generates accurate approximations for optimal solutions to RL problems without sacrificing computational efficiency. Furthermore, the model is able to efficiently update its representations in response to environmental changes or new objectives. Critically, this update mechanism demonstrates stronger alignment with observed human and animal behavior in replanning scenarios. Additionally, because SR-IS shares the same theoretical foundation as linear RL, specifically that computational costs are incurred when deviating from the default policy, it offers the same unified framework for understanding inflexibilities in decision making, including habits, Pavlovian-instrumental transfer effects ^49–51^, and constraints on cognitive control ^52–56^.

Recent work building upon the linear RL framework, proposed that cognitive maps could be learned through compositional positioning of object and barrier representations within an open-space baseline map ^57^. While SR-IS offers an alternative approach by creating maps purely through learning, these models can coexist complementarily - one explaining compositional map building and the other addressing learning-based map construction. Importantly, many situations, including the human revaluation tasks simulated in the current study, cannot be handled through compositional positioning of familiar barriers and objects. Thus, while the compositional model offers valuable insights, it cannot explain the specific behavioral patterns in revaluation tasks that are successfully captured by SR-IS.

SR-IS leverages a fundamental computational advantage of the linear RL framework: the low-rank matrix update. Because it learns the DR, SR-IS can exploit this method to adapt rapidly when environments change, such as when barriers are added or goals relocated. This update inverts only a small matrix for the altered transitions rather than recomputing the entire DR, achieving substantial computational savings while remaining exact. Since SR-IS produces an unbiased DR estimate via importance-sampled TD learning, these updates work much like in the Complete linear RL, combining computational efficiency with online learning. However, this approach generates a testable prediction such that replanning quality should depend on how well the baseline representation is learned. If the DR is only partially learned, the low-rank update, designed for exact matrices, may fail to produce correct adaptations. This could be tested by asking whether humans show degraded replanning when structural changes occur after limited experience. More broadly, this suggests that rapid replanning methods may require extensive prior learning ^58^, predicting quick, accurate replanning in familiar environments but degraded performance when changes occur before sufficient exploration.

Our current work uses a TD learning rule where the agent updates its predictions based only on immediate experiences. Recent modeling work, however, suggests that human learning may be better captured by models that incorporate eligibility traces, a mechanism that maintains memory of recently visited states and updates them based on later outcomes. Kahn et al. ^59^ showed that learning rules with strong eligibility traces provide better fits to human behavioral data than those using only immediate updates. In principle, the SR-IS framework can accommodate eligibility traces. The key insight is that any learning rule that works for standard SR can also work for learning the DR. Moreover, importance sampling allows us to learn it while behaving according to current goals by weighting our learning updates appropriately to account for the difference between how we are behaving and the default-dependent representation we are trying to learn. Future work could explore whether incorporating eligibility traces into SR-IS provides a better account of neural and behavioral data than the version presented here. This would test whether the human brain uses eligibility traces not just for learning about current behaviors, but also for maintaining flexible, reusable representations of environment structure.

There is a long history of using importance sampling in RL, particularly for off-policy learning. In the typical setting, it is applied in offline RL to learn an optimal policy from data generated by an exploratory policy ^60–63^. Much prior work has focused on reducing the high variance of the importance sampling estimator ^64–68^. Our application in SR-IS, however, inverts this usual use case: instead of learning specialized policies from general experience, it uses importance sampling to learn general representations from specialized behavior. This approach connects most directly to Todorov’s work on linearly solvable Markov decision processes ^36^, the theoretical foundation for the linear RL model, where importance sampling enables model-free learning of value functions. By repurposing importance sampling from a tool for policy optimization to one for representation learning, SR-IS shows how this technique can offer new insights into biological planning and cognitive flexibility.

Despite its advantages, SR-IS operates under specific constraints. The model requires that the decision policy maintains non-zero probability for all actions that the default policy might take, it cannot function with purely greedy policies where one action receives probability one and all others zero. This constraint suggests that biological systems may maintain behavioral variability not simply as exploration, but as a computational necessity for flexible representation learning. While assigning near-zero probabilities to certain actions creates the variance that explains human behavioral patterns in our analyses, extreme probability imbalances could make the model impractical in large-scale environments. Future studies should systematically test SR-IS in larger, more realistic settings to determine whether biological systems employ similar mechanisms and, if so, how they manage these constraints.

This computational need for variability offers a new perspective on stochastic behavior in planning. Rather than treating randomness as a byproduct of information-seeking ^69;70^ or uncertainty-driven learning ^71–74^, SRIS frames behavioral variability as essential for maintaining reusable representations. This view leads to a specific, testable prediction where variability should correlate with successful adaptation, especially in novel environments, even when it provides no new information about rewards or task structure. This stands in contrast to information-theoretic accounts of exploration, where variability is valuable only insofar as it reveals unknown contingencies ^69;70^. Under SR-IS, an agent that maintains moderate behavioral variability should transfer more effectively to new goals than one following near-deterministic policies ^75^, even in fully observable, deterministic settings where extra information offers no benefit.

Another limitation of the SR-IS model, carried over from the linear RL framework, is its assumption of fully controllable transition dynamics, which holds in environments like mazes but not in many biologically relevant, stochastically controlled tasks. While we have previously extended the linear RL framework to handle stochastic tasks through an approximation (first solving for optimal dynamics as if transitions were deterministic, then projecting this solution onto achievable policies), this approximation may not always hold ^37^. There are scenarios in which this approach fails, when there is a mismatch between controllable and achievable transitions leading to systematic errors. These arise when stochasticity in action outcomes significantly alters the value of future states, such that assuming full controllability misleads planning at other, predecessor steps. This generally leads to an optimistic bias. Whether humans show these optimistic biases remains to be tested. Humans may well exhibit SR-IS-like biases in some situations, but unlike our model which would always fail under these conditions, humans appear capable of overcoming these biases at least some of the time. This flexibility, absent in SR-IS, likely reflects their ability to detect when simplified strategies fail and revert to more expensive MB planning, resulting in either reduced performance or systematically slower responses in challenging scenarios.

The control cost parameter in the linear RL framework, inherited by SR-IS, reveals important connections between computational constraints and behavioral flexibility. This parameter governs the trade-off between optimal performance and adherence to default behavior where high control costs produce rigid, default-dominated behavior, while low costs enable more optimal but computationally demanding policies. Critically, approaching zero control cost, while theoretically yielding optimal behavior, creates numerical instability as the framework must multiply small probability values by exponentially large reward terms. This suggests connections to known neural limitations such as maximum firing rates, precision bounds, and rational models under representational noise ^76–78^. Our current implementation assumes fixed control costs that vary across individuals, potentially explaining individual differences in cognitive flexibility versus habitual responding. However, dynamic adjustment of control costs within tasks ^44^, allowing flexible shifts between exploratory and exploitative modes, remains unexplored in our framework. Such dynamic control could explain moment-to-moment variations in behavioral variability and performance, connecting our model to broader questions about cognitive control, effort allocation, and the neural implementation of gain modulation.

Beyond cognitive science, our approach could guide the development of novel artificial intelligence systems that can plan efficiently and adapt to new situations. The SR has already influenced several advances in artificial intelligence, including the development of more generalized environmental features known as “successor features” ^79^) and methods for sub-goal discovery ^80;81^. These applications leverage the SR’s ability to create environmental representations that facilitate transfer learning, allowing systems to apply knowledge from one task to enhance performance on others ^82^. We propose that incorporating importance sampling could provide a straightforward yet powerful enhancement to these approaches. By addressing bias in the SR, importance sampling should yield more generalized features and more effective sub-goal identification, while maintaining computational efficiency.

## Methods

### Reinforcement learning

RL addresses planning in environments characterized by states, actions, and rewards. The goal is to learn value functions that predict expected future rewards. The value of a state under a specific policy is the expected cumulative reward when following that policy, while the optimal value function represents the maximum achievable value for that state across all possible actions. The challenge of RL can fundamentally be understood as learning to accurately predict these value functions. Two distinct approaches have been developed to tackle this challenge: Model-based algorithms explicitly learn the environment’s dynamics by learning the transition probability between states and the reward function. The second approach is model-free RL, which directly updates a cached value of the estimate itself using algorithms such as TD-learning ^26^. Specifically, after observing a transition *s*→*s*′ with a reward, *r*(*s*), TD calculates a reward prediction error, *δ*, to update the value of state *s, V* (*s*).

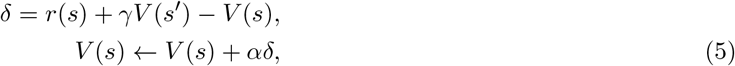

where *γ* is the discount factor parameter, *α* is the learning rate parameter which controls the step size of the agent as it updates its estimate of the value function.

### Successor representation

The SR, defined in Equations 1-2, has an important matrix formulation as shown by Russek et al ^24^. Let **T**^*π*^ be the one-step transition matrix under policy *π*, where element 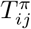 represents the probability of transitioning from state *i* to state *j*. This transition matrix can be written as:

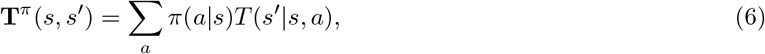

where *T* (*s*′ | *s, a*) is the transition probability given action *a*. Using this transition matrix, the SR can be expressed as:

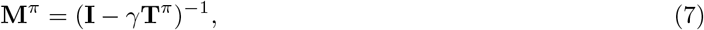

where **I** is the identity matrix and *γ* is the discount factor. Values are then computed as **v** = **M**^*π*^**r**, where **v** is the vector of values across all states, and **r** is the reward vector.

### Linear RL

The linear RL model ^37^ provides a closed-form approximation to the optimal value function for a specific class of finite-horizon Markov decision problems with deterministic transitions, such as maze navigation toward a specific goal. At its core, the model optimizes behavior by maximizing a gain function that balances rewards against a control cost term. The gain function *g* for state *s* is defined as:

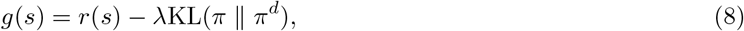

where *r*(*s*) represents the reward at state *s*, and *λ >* 0 is a control cost parameter. The second term measures the Kullback-Leibler divergence between two policies: the agent’s decision policy *π* and a default policy *π*^*d*^. This divergence is zero only when the policies match exactly (*π* = *π*^*d*^) and is positive otherwise. Throughout our simulations, we use a uniform default policy where all transitions to neighboring states are equally likely (though the method works regardless of this assumption). Such a uniform default policy introduces stochasticity into decision-making without biasing the decision policy toward any specific actions. It can be shown that the optimal value function for this problem has an analytical solution, which can be viewed as an approximation to the value function of the original RL problem without control costs. To derive this solution, we first define the one-step state transition matrix under the default policy. Since we assume the default policy is uniform throughout this paper, this matrix is equal to **T**, where each element *T*_*ij*_ represents the probability of transitioning uniformly from state *i* to state *j*. For brevity, and to keep the notation uncluttered, we omit the explicit dependence on the default policy, as we assume it to be uniform throughout our analysis.

Partitioning the states into nonterminal and terminal (goal) states, and defining **t** = **T**_*NT*_ exp(**r**_*T*_ */λ*), where **T**_*NT*_ is the transition matrix from nonterminal to terminal states and **r**_*T*_ is the vector of rewards for each terminal state, it can be shown that optimal values across all nonterminal states is given by:

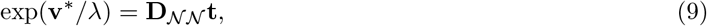

where **v**^*\**^ is the vector of optimal values across all nonterminal states, and **D**_*NN*_ is the submatrix of the DR matrix **D** corresponding to nonterminal states. The DR matrix is defined as:

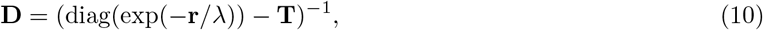

where **r** is the reward vector across all states (assuming non-zero rewards at terminal states). If we assume that the reward at all nonterminal states is equal to *r*_*N*_, then we can write **D**_*NN*_ = *γ***M**_*NN*_, where we have defined *γ* = exp(*r*_*N*_ */λ*). Here, **M**_*NN*_ is the submatrix of **M** corresponding to nonterminal states, defined as:

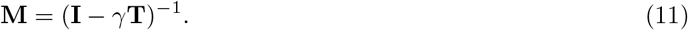

Throughout the paper, we generally refer to both **D** and **M** as the DR matrix because they are simply related to each other through the constant *γ*, except where this might cause confusion. The DR matrix, defined in Equation 11, shares a strong similarity with the SR matrix (Equation 7) - in fact, the DR matrix can be understood as the SR under the default policy. Moreover, *γ* can be interpreted as equivalent to the discount factor in Equation 7. It is important to note, however, that although the DR (Equation 11) and SR (Equation 7) are structurally similar, they are fundamentally different: the DR matrix is based on the transition structure **T**, whereas the SR matrix is based on the policy-dependent transition structure **T**^*π*^, defined with respect to the agent’s learned decision policy *π*.

### Decision policy

The decision policy is defined using a softmax rule:

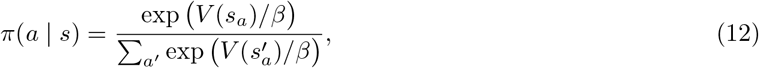

where *s*_*a*_ is the successor state associated with action *a*, and *β* is the temperature parameter indicating the degree of exploration (i.e., randomness). Here, *V* (*s*) is the value function for state *s* computed by either algorithm. In all our analyses, we assumed *β* = *λ* for the SR-IS algorithm.

### Replanning

A key advantage of the linear RL framework and its DR matrix is that it enables efficient low-rank updates to the predictive map when the environment changes. This contrasts with both classic model-based approaches and the SR, which typically require recalculating the entire map even for minor environmental changes.

Formally, when an environmental change modifies matrix **L**_0_ = diag(exp(−**r***/λ*)) −**T** to a new matrix **L** (due to alterations in either **T, r**, or both), we can apply the Woodbury matrix inversion lemma ^41;42^. Given 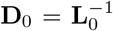, this lemma allows us to compute the new matrix **D** = **L**^*−*1^ through a computationally efficient low-rank update of **D**_0_.

Now, let us introduce two sparse matrices **R** and **C** of size *J* × *S* and *S* × *J* respectively, where *S* is the total number of states and *J* is the number of states for which **L**_0_ has changed. Let **j** = (*j*_1_, *j*_2_, …, *j*_*J*_) be a vector of length *J* containing the indices of all those states, where *j*_*i*_ denotes the *i*-th element of the vector. For each changed state, matrix **R** contains rows representing the difference between the corresponding rows in **L** and **L**_0_, such that **R** = **L**(**j**, :) − **L**_0_(**j**, :). Matrix **C** acts as a selection matrix: for each changed state *j*_*i*_, it contains a column with a value of 1 in row *j*_*i*_ and zeros elsewhere. Using these matrices **C** and **R**, we can efficiently express the change in **L**_0_ as:

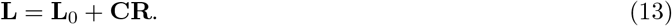

Using this formulation, we can apply the Woodbury matrix inversion lemma ^41;42^ to express **D** (the inverse of as a low-rank update of **D**_0_:

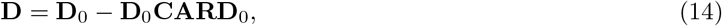

where **A** = (**I** + **RD**_0_**C**)^*−*1^. While this formulation still requires matrix inversion for replanning, it operates on a much smaller matrix: **A** has dimensions *J* × *J*, where *J* is the number of changed states and is typically much smaller than the total number of states *S*.

Since SR-IS learns the DR, we can use these equations for replanning with the SR-IS model. In this setting, we use the SR-IS algorithm to learn **D**_0_ and apply Equation 14 to update the matrix if a new barrier or goal is introduced. In practice, baseline map imperfections occasionally produce negative z-values (where **z** = **D**_*NN*_ **t**) for some states. However, given Equation 9, **z** = exp(**v**^*\**^*/λ*), and should therefore always be positive. To address this, we offset all z-values by the minimum value, ensuring they remained positive as theoretically required. This correction worked efficiently in our simulations, though it may not be the most accurate approach and could be refined in future work.

### Generalization of the SR-IS algorithm

We derived the SR-IS algorithm (Equations 3 and 11) under the assumption that all nonterminal states share the same reward. However, it is straightforward to extend the algorithm to account for unequal rewards across nonterminal states. Specifically, define ***γ*** = exp(**r**_*N*_ */λ*), where **r**_*N*_ is the vector of rewards across nonterminal states. We then have:

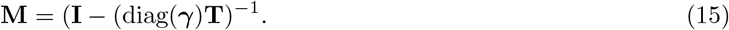

Consequently, the following learning rule should be used:

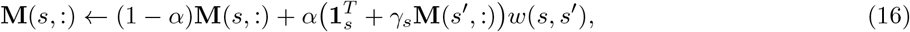

where *γ*_*s*_ is the *s*th element of ***γ***.

An important feature of the linear RL framework inherited by SR-IS is the relationship between the discount factor *γ* and control cost parameter *λ*. Since ***γ*** = exp(**r**_*N*_ */λ*), these parameters are inversely related: lower *γ* corresponds to higher *λ*, which increases the cost of deviating from the default policy. This creates a principled trade-off where higher control costs produce behavior closer to the default policy (promoting exploration and stochasticity), while lower control costs permit greater deviation to optimize rewards (enabling exploitation). In SR-IS, this means the same parameter that determines temporal discounting in the learned representation also regulates how much the agent’s behavior can diverge from the default policy, creating a natural coupling between representation learning and behavioral expression.

### Simulation and fitting procedure

#### Fixed parameters across simulations

All of the maze simulations used a custom maze environment created using OpenAI’s gym ^83^. For all of the simulations we assumed a uniform default policy. Across all our simulations, unless otherwise specified, we set parameters *β* and *λ* to be 1, the learning rate (*α*) to be 0.05, the reward at non-terminal states (*r*_*N*_) to be − 0.1, and the reward at terminal states to be 10. The SR model had the same parameters except we defined the reward at non-terminal states to be 0 and its discount factor to be 0.9 (equal to the SR-IS *γ*). We define a “step” as when the agent takes an action, receives a reward, and consequently updates its representation. We vary the steps on each simulation based on how many was needed to achieve a low standard error of the mean. For every model that has a representation of the environment we assume a (state, state) representation. All of the models employ the softmax function (Equation 12) to make decisions. Across all tree-based revaluation analyses, we used auxiliary terminal states connected to each goal state.

#### Convergence and replanning

For the convergence simulations in Figure 2**a**,**b** as well as the revaluation problem in Figure 2**a**,**c** a 7 × 7 four-room maze was considered. For the convergence simulations, we set the terminal state to the bottom-right corner of the maze. We iterated for 1.5k episodes and took the mean error between the agent’s representation and the complete DR every 10 steps. We used episodes (sequences from initial to terminal state) instead of steps for this simulation only to make the figure easier to read. For the revaluation simulations, we trained the SR and SR-IS agents with the assumption that every corner (*r*_1*−*9_) was a terminal state. During training we set the payout of the terminal state in the top-right to have a reward of 10, setting all the others to have the same reward as non-terminal states. At test, we set the reward of the terminal state we trained towards to be the same reward as non-terminal states and alternated making the reward of every other terminal state 10. Depending on which room these terminal states were in we classified these as “Same Room” and “Different Room” trials. We trained and evaluated the models across 50 runs each consisting of 15k steps.

For the simulations presented in Figure 2**d**-**f**, a 10 × 10 maze was used. For both settings, the initial goal state was set to be in the bottom right corner of the maze. After initially calculating the DR for the first terminal state, we moved the terminal state to every other open state and re-used the initial DR to solve for an updated DR using Equation 14. We evaluated all the models across 10 simulations. It is important to note that unlike the analysis done by Piray and Daw ^37^, which computed a goal-independent representation, we require a goal state to test SR-IS as with no goal state the learning policy would be random and we would not need importance sampling.

#### Policy revaluation from Russek et al. ^24^

For the simulations presented in Figure 3**b**-**d**, a 11 × 11 maze was used. Initially we set the payout of r_1_ to be 10 and r_2_ to be 0 and we trained the model(s) for 40k steps on this payout structure. At test, we changed the reward of r_2_ to be 20 and used the representations learned during training to generate a new policy based on the new reward payout.

#### Simulation of Momennejad et al. ^25^

For the simulations of experiments by Momennejad et al. ^25^ in Figure 4**a**-**e**, we followed the payout structure from the paper for the terminal states. We trained the agents in a similar fashion to how the participants experienced the task where they started 14 times from state 1, 7 times from states 2 and 3, and 2 times from each terminal state. Because the linear RL framework (and SR-IS) requires terminal states to be pre-specified, whereas SR (and other online algorithms) does not, the two are not directly comparable. To resolve this, we introduced auxiliary terminal states connected to each goal state, placing the DR and SR on equal footing. This adjustment was used across all analyses comparing SR-IS with SR in tree-based tasks testing revaluation effects. It reflects the idea that agents do not know which states are terminal until they are encountered. We set *λ* = *β* = 5 to avoid overflow of the exponential due to large reward values in the decision policy, and averaged results across 800 runs at test time.

#### Kahn and Daw ^59^

For the simulation in Figure 5**d**-**i**, we used the source code provided by Kahn and Daw ^59^. To simulate the SR-IS agent we modified their code for the SR agent by adding an importance sampling term outlined in Equation 3 to the TD update for the SR agent and extending the backbone to use linear RL framework. The analysis we show is an extension of their original analysis which fitted simulated model decisions, after an observation trial, to a linear mixed model with the coefficients designed in such a way to tease apart the effects of SR vs. model-based decision making strategies. We use all of the same sampling strategies that they did for the SR as they share the same variables, with the exception of defining a range for *λ* to be sampled uniformly between (0.01, 0.2). Note that, as in the original study, this task also introduces auxiliary terminal states connected to each goal state. Please refer to the section above for further rationale.

For model fitting and comparison, we considered four models. The SR and SR-IS models were implemented as described in Equations 9–12. The model-based agent was given perfect knowledge of the transition structure of the environment, *T* (*s*′ | *s, a*), and computed values at each timestep through iterative maximization such that *V* (*s*) = max_*a∈A*_ *V* (*s*_*a*_), where the value of a state is defined as the maximum over the values of its successor states. The hybrid SR+MB agent was constructed as a linear combination of the SR and model-based accounts in value space, following the original study ^44^. For all models, we assumed that the reward mean for terminal states was learned via a delta rule with *α*_r_ as the reward learning rate. Model parameters were estimated using standard fitting procedures that we and others have used previously ^46;84;85^. The parameters fit for each model were: MB(*β*), MF(*β*), SR(*β*), Hybrid SR+MB(*β*_1_, *β*_2_), SR-IS(*γ, λ, β*). Specifically, parameters were fit separately for each subject using a maximum-a-posteriori procedure with Gaussian priors. Appropriate transformations were applied to each parameter, including a sigmoid transform for parameters bounded between zero and one and an exponential transform for parameters constrained to be positive. Gaussian priors were assumed to have zero mean and variances sufficiently large to allow parameters to vary over a wide range with minimal influence of the prior (100 for temperature parameters and 6.25 for others, consistent with prior work and our previous modeling studies ^85–87^). Model evidence for each subject across all candidates was quantified using the Laplace approximation and subsequently subjected to a random-effects Bayesian model selection to identify the best-fitting model at the group level ^45;88;89^, implemented using the cbm package in python ^46^.

#### de Cothi et al. ^47^

For the simulations in Figure 6**a**–**d**, we used the MATLAB source code provided by de Cothi et al. ^47^. To simulate the SR-IS agent, we modified their SR agent code by adding the linear RL machinery for policy computation and the importance-sampling term outlined in Equation 3. We fit the SR-IS model using the fitting procedure from the original paper (MATLAB’s fmincon). The parameters fit for each model were: MB(*γ*), MF(*α, γ*), SR(*α, γ*), Hybrid SR+MB(*α, γ, w*), SR-IS(*α, γ, λ*). We also used the plotting code provided in the original paper to reproduce the model comparison across trials and maze configurations, ensuring direct comparability with the original results. We then performed random-effects Bayesian model selection, with model evidence approximated using the Bayesian information criterion (BIC) computed separately for each model and subject, implemented using the cbm package in MATLAB ^46^.

#### Stochastic two-step task

For the simulations in Figure 7**a**-**e** an environment with 7 states was used. The first state (*S*_1_) transitioned to two successor states deterministically (*S*_2_ and *S*_3_) with actions *A*_1_ and *A*_2_ respectively. Once in states *S*_2_ and *S*_3_ the agent again had the choice of two actions (*A*_1_ and *A*_2_) with each leading to a terminal state with some reward. In *S*_3_ the actions deterministically led to the same reward of +0 and +1. However, in *S*_2_ the actions were stochastic with *A*_1_ and *A*_2_ leading to two different states with rewards of −10 and +10 (with 50-50% chance). Thus, the expected value of *S*_3_ is higher than *S*_2_. At revaluation we changed the reward structure of the terminal states to make the expected value of *S*_2_ higher than *S*_3_. We trained the agents for 250 steps and averaged across 500 different runs.

## Data availability

No new data were collected for this study. Simulated data are provided along with the code.

## Code availability

Analyses were performed using custom code developed in Python, MATLAB, and Julia. The complete code repository is available at https://github.com/ArminBaz/SR-IS. We utilized the original code repositories from de Cothi et al. ^47^ and Kahn and Daw ^44^ to extend their respective analyses. Model fitting and comparison were conducted using the cbm package, which is available at https://github.com/payampiray/cbm_python.

