## Supplementary figures for "Efficient Learning of Predictive Maps for Flexible Planning"

### Supplementary Materials:
#### Efficient Learning of Predictive Maps for Flexible Planning

Armin Bazarjani[1*] and Payam Piray[1*]

[1]Department of Psychology, University of Southern California, Los Angeles, CA

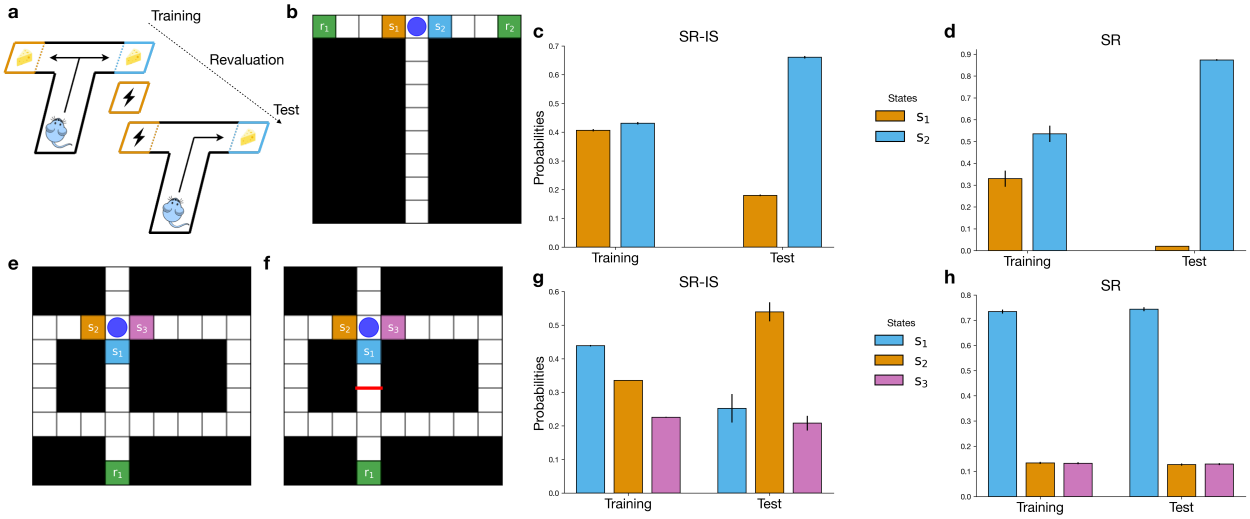


Supplementary Figure 1: **Performance of SR-IS on Tolman's tasks.** (**a-d**) Latent learning task. (**a**) The task structure. Rats were initially allowed to forage freely in a maze with two rewarding end boxes where they would receive an equal amount of reward in either end box. In the next phase, one of the boxes resulted in a shock without warning, devaluing that terminal state. (**b**) The maze used to simulate this task, the blue circle represents the agent's current location, the green squares represents the reward location, the orange and blue squares represent the states available for transition to the agent $s_1$ and and $s_2$ respectively. (**c**) SR-IS agent is able to change it's preference from 50-50 towards the more rewarding state. (**d**) SR agent is also able to change its preference. (**e-h**) Detour task. (**e**) The environment initially has three possible paths, with shortest being to go straight through state $s_1$. (**f**) In the next phase, the path going straight is blocked and the agent must be able to update its learned representation to account for the barrier, ideally choosing the next shortest path through state $s_2$. Again the blue circle and the green square represent the agent's location and the reward state respectively. The green, orange, and purple squares represent the states available for transition at the junction ($s_{1-3}$). (**g**) SR-IS is able to correctly update its representation and select the path that avoids the barrier. (**h**) SR is unable to account for the change in the transition structure of the environment and still prefers the, now blocked, shortest path. The reward of non-terminal states in the detour task was set to -0.01 (making $\gamma = 0.99$ instead of 0.9), all other parameters were kept the same as the default set of parameters reported in the main text. Error bars represent the standard error of the mean.

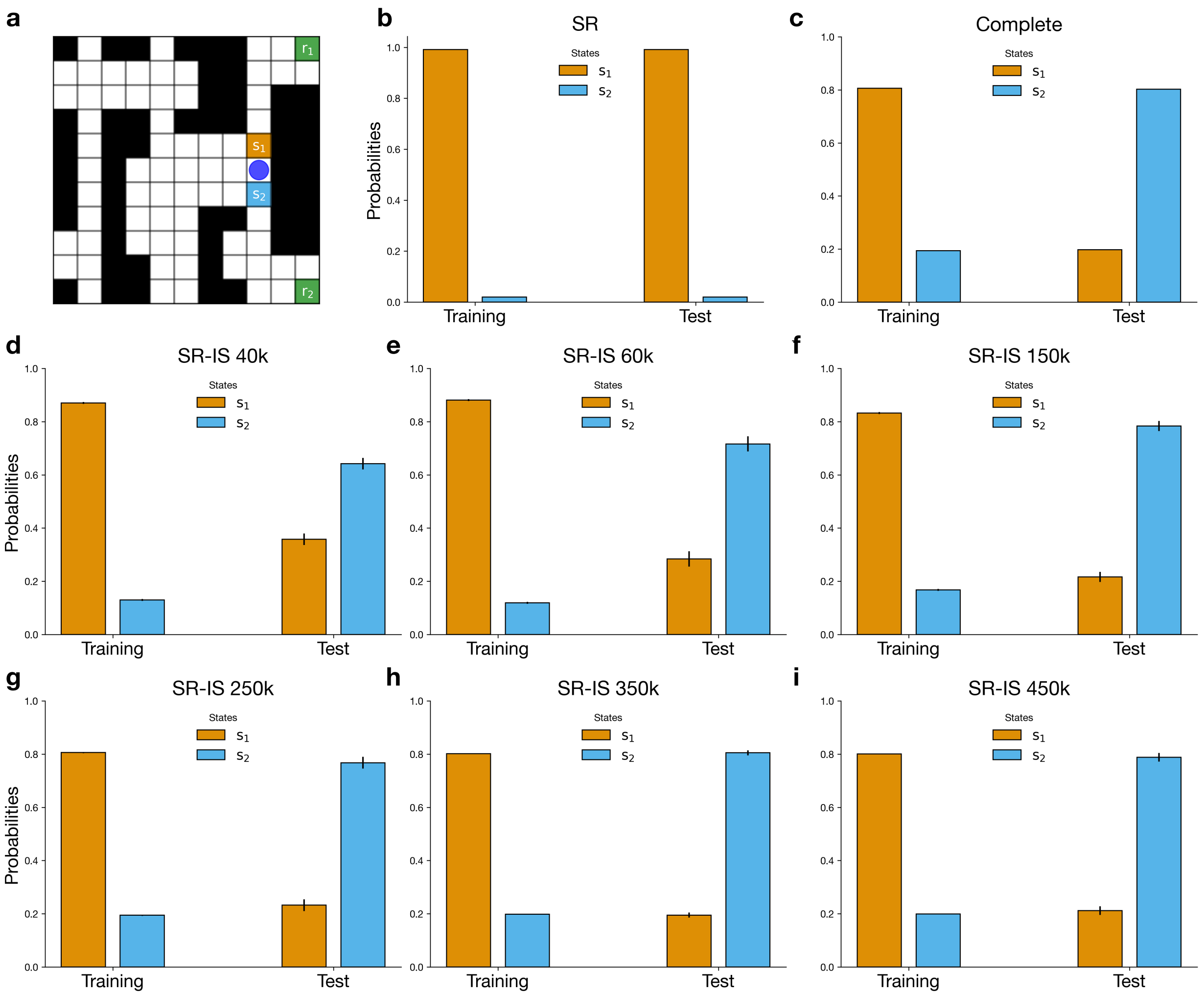


Supplementary Figure 2: **Performance of SR-IS on policy revaluation task from Russek et al.[1].** (**a**) The maze environment, the agent's location is shown with a blue circle and two neighboring states ($s_1$ & $s_2$) showing different paths to the two opposing terminal states ($r_1$ & $r_2$). (**b-i**) We show the probability of each agent in selecting either state $s_1$ or $s_2$, demonstrating the agent's preferences over both terminal reward states $r_1$ and $r_2$ during training and testing. (**b**) SR (SR-MB) agent cannot properly adapt it's value representation to accommodate the change in reward. (**c**) The complete (linear RL) model is able to perfectly switch its preference when the reward changes due to its complete knowledge of the environment. (**d-i**) Performance of SR-IS with different training lengths, measured through number of steps. With more training, the bias exhibited by SR-IS becomes less salient. Error bars represent the standard error of the mean.

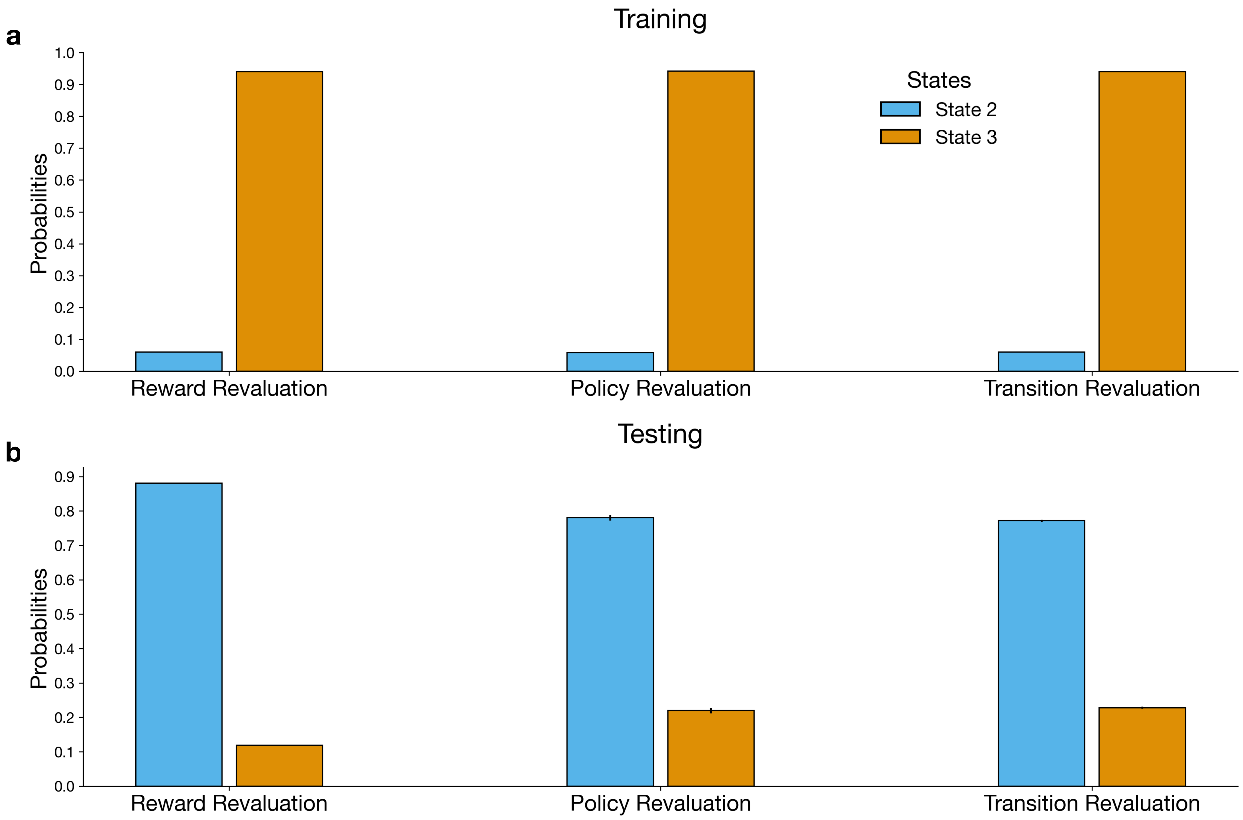


Supplementary Figure 3: **Performance of SR-IS model on Momennejad et al.'s[2] replanning tasks.** (**a**,**b**) Probability of SR-IS selecting between states 1 and 2 both during training and at test. The agent was in the same fashion that the human participants were exposed to the task. Error bars (might not be visible) represent the standard error of the mean.

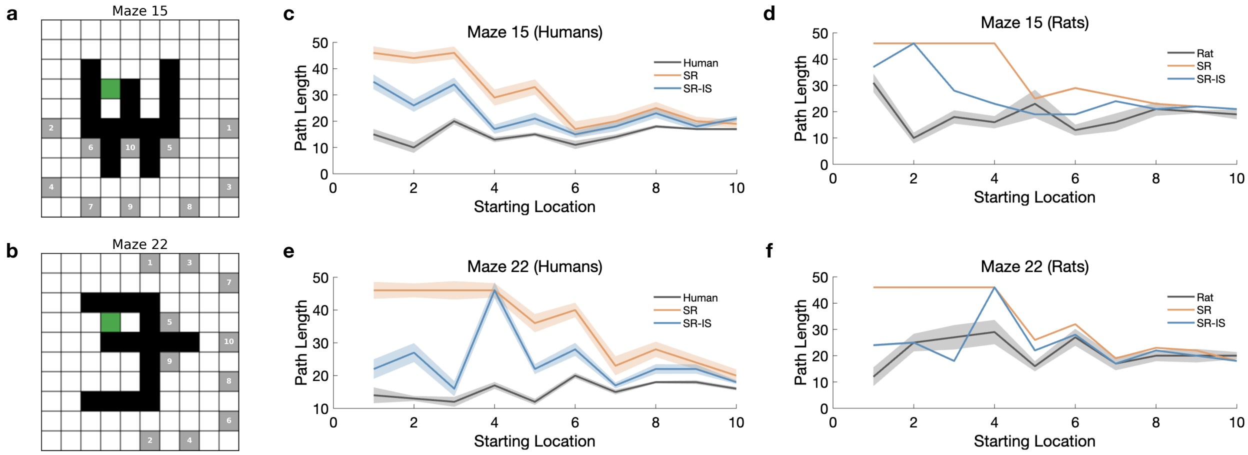


Supplementary Figure 4: **SR-IS path length is closer to that of humans and rats in de Cothi et al.[3].** (**a**, **b**) Two distinctive examples selected from a total set of 25 mazes, each demonstrating the challenge of policy dependence that arises when navigating from different starting states. (**c-f**) For both maze 15 and maze 22, the SR-IS model demonstrates superior performance by providing a closer match to human and rat path lengths from every starting location. Error shading represent the standard error of the mean.

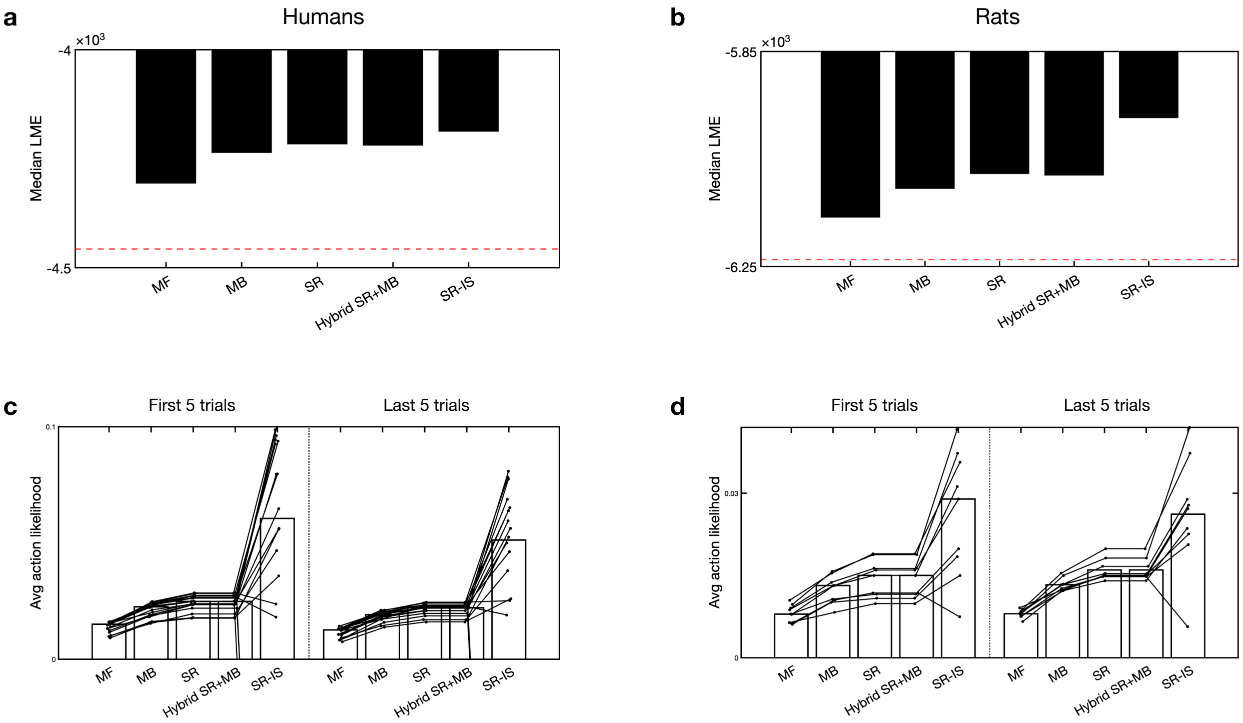


Supplementary Figure 5: **Maximum likelihood analysis across human and rat trajectories** (**a**, **b**) Median, log model evidence (LME) across both humans (**a**) and rats (**b**). The value estimates generated by the SR-IS agent provide a more likely explanation of the biological behavior than either the MF, MB, SR, or Hybrid SR+MB agents. (**c-f**) By outputting the average action likelihood per timestep we can see that this trend is true across all individuals in both humans (**c**) and rats (**d**) and robust throughout exposure to a maze configuration. Indicating SR-IS is the best fitting model from start to finish across all the mazes.
